# Speed limits of protein assembly with reversible membrane localization

**DOI:** 10.1101/2021.01.29.428888

**Authors:** Bhavya Mishra, Margaret E. Johnson

## Abstract

Self-assembly is often studied in a three-dimensional (3D) solution, but a significant fraction of binding events involve proteins that can reversibly bind and diffuse along a two-dimensional (2D) surface. In a recent study, we quantified how proteins can exploit the reduced dimension of the membrane to trigger complex formation. Here, we derive a single expression for the characteristic timescale of this multi-step assembly process, where the change in dimensionality renders rates and concentrations effectively time-dependent. We find that proteins can accelerate complex formation due to an increase in relative concentration, driving more frequent collisions which often wins out over slow-downs due to diffusion. Our model contains two protein populations that associate with one another and use a distinct site to bind membrane lipids, creating a complex reaction network. However, by identifying two major rate-limiting pathways to reach an equilibrium steady-state, we derive an accurate approximation for the mean first passage time when lipids are in abundant supply. Our theory highlights how the ‘sticking rate’, or effective adsorption coefficient of the membrane is central in controlling timescales. We also derive a corrected localization rate to quantify how the geometry of the system and diffusion can reduce rates of localization. We validate and test our results using kinetic and reaction-diffusion simulations. Our results establish how the speed of key assembly steps can shift by orders-of-magnitude when membrane localization is possible, which is critical to understanding mechanisms used in cells.

## I. Introduction

The speed with which populations of interacting biomolecules such as proteins relax to an equilibrium steady-state is an important reference point for understanding how fast they can self-assemble in the cell. In clathrin-mediated endocytosis, the construction of a protein-coated membrane vesicles takes ~60s, across a variety of cell types^1^. However, the ultra-fast form of endocytosis is over 1000 times faster, while still requiring assembly of proteins and budding of the membrane^2^. The physical variables of protein concentrations, membrane lipid composition, association rates, diffusion, and cell geometry that determine these timescales thus must support this similar function at quite divergent timescales. Experiments performed *in vitro* can determine timescales of association and assembly under specific conditions, but transitioning to the cell requires, at the very minimum, accounting for changes in concentrations, membrane composition, and system dimensions. We focus here on how the ability of protein populations to reversibly localize to membranes, an essential step in endocytosis, in signal activation^3^, in cell polarity establishment^4^, and in cell adhesion^5^, will alter timescales to complex formation from an initially unbound population. By deriving characteristic timescales for this process, which could be directly mapped to an *in vitro* system, we can predict how transitions in environment and components could accelerate or decelerate key steps in assembly processes on membranes.

In complex biochemical reaction networks in chemistry and biology, bimolecular association represents a fundamental building block, and thus provides a well-studied theoretical foundation for our model. Kinetics of the well-mixed rate equations are analytical soluble. The kinetics of explicitly spatial models of reaction kinetics are slower than rate-equations as equilibrium is approached^6, 7^, but in 3D, the time-dependence is otherwise accurately predicted by rate equations. In dimensions less than three, rate-equations fail to fully capture the effects of diffusional dynamics on kinetics at all time-scales^8^, but optimal macroscopic rates can be derived to approximate the bulk reaction kinetics^9–11^. Characterizing how bimolecular association depends on kinetic and geometric or spatial parameters is simplified by the definition of characteristic times, such as the first passage time (FPT)^12^ or mean first passage time (MFPT)^13, 14^. While FPTs describe the time for a stochastic process to reach some specified threshold, such as the time to reach a target in confinement^15, 16^, the MFPT represents a population average, and is thus descriptive of both deterministic and stochastic models. For example, the MFPT of a first order process with rate *k* is simply given by 1/*k*, which is ln (2)^−1^~1.44 times longer than the half-time (50% completion). For bimolecular association, the MFPT must also depend on initial concentration of species, but the reaction rate constants remain a primary determinant of characteristic times. Although the association rate for a pair of biomolecules fundamentally depends on electrostatics and molecular structure, for example^17^, we will assume this contribution to the rate is known. Using the Smoluchowski and Collins-Kimball model^18^, we will explicitly account for the additional dependence of the rate on diffusion and spatial dimension. This is important, because incorporating the effects of geometry and diffusion on reaction rate constants allows us to predict timescales from the coupled ordinary differential equations (ODE), which is significantly simpler than partial differential equations (PDE). Along with the concentrations, the reaction rates can effectively predict either the MFPT or half-time in two-component systems.

In complex biochemical reaction networks, however, predicting kinetics and relaxation times when more than two species are coupled together nonlinearly becomes rapidly intractable, often necessitating numerical solutions or prior knowledge of the rate-limiting steps^19^. In our model, we have added one additional component and 9 additional reaction channels, due to the additional ‘domain’ of the membrane, extending significantly beyond simple bimolecular association but with constraints on binding rates as they change from 3D to 2D. Diffusion can play a critical role via its reduction from 3D to 2D (~100 fold), and its impact on localization times to the surface. We note that diffusion-influenced bimolecular reactions have also been theoretically studied when additional complexity is added not via additional reactants, but via multiple sites per reactant^20, 21^, or via tethering between sites^22, 23^. For these more complex reaction networks, solutions can always be found numerically using non-spatial rate-equations or spatial reaction-diffusion methods^24^, both of which we use here for validation. Stochastic single-particle reaction-diffusion methods in particular are ideal for quantifying the role of diffusion and density fluctuations in kinetics of bimolecular association^9^, multi-site phosphorylation^25^, or oscillators^26^, for example. Although dramatic new behavior can emerge in spatial representations, for simpler networks like the one studied here, non-spatial solutions tend to be qualitatively (often quantitatively) similar^27^.

Bimolecular association with the possibility of dimensional reduction to a surface represents a more complicated but still fundamental building block of many cell-biology processes; understanding the origins of its characteristic timescales is important for assessing speeds in yet more complicated systems. By allowing transitions between the surface, the bimolecular association rates and the protein concentrations of the two-component system can be thought of as effectively time-dependent. The role of dimensional reduction is of long-standing interest in biology; Adam and Delbruck quantified when a ligand can more efficiently locate its membrane-bound receptor target by adsorbing to the membrane and searching in 2D^28^. Berg and Purcell predicted how receptor density and affinity, along with diffusion, can determine activation times of receptors^29^. More recently, the MFPT for a molecule to reach a surface target was shown to be sensitive to the switching time from solution to surface^30^. In the nucleus, dimensional reduction to the approximately 1D DNA was predicted to help proteins locate promoters^31^, which has been observed experimentally^32^. In all of these examples, one of the ‘reactants’ is already localized to the lower-dimensional domain, and thus the predicted time-scales do not describe how populations of interacting molecules can exploit dimensional reduction to control assembly speeds.

Dimensional reduction impacts not only the kinetics of association reactions, but also the probability of proteins partitioning to bound versus unbound states (Fig 1a). For receptor targeting, this means scaffold proteins that also are lipid bound are significantly more likely to be bound to their receptor^33^, as has also been observed experimentally^34^. The effective increase in concentration on the membrane can cause increases in formation of multi-protein oligomers^35^, and in signaling networks, it can promote bistability^3^. In a recent study^36^, we developed the model we use here to derive an effective equilibrium constant that predicts how concentrations, binding affinities in 3D and 2D, and volume to area ratios will alter the equilibrium. We found dimensional reduction triggered dramatic increases in complex formation for physiologic regimes. This framework also extends to proteins that do not bind directly to the membrane themselves, but can form a scaffold on the surface via multi-valent interactions^36^, as has also been observed experimentally for clathrin assembly on *in vitro* membranes^37^. Using reaction-diffusion simulations, this work tested how dimensional reduction impacted the speed of assembly for physiologic binding pairs (Fig 1), and here we develop a predictive analytical theory.

**Figure 1.**
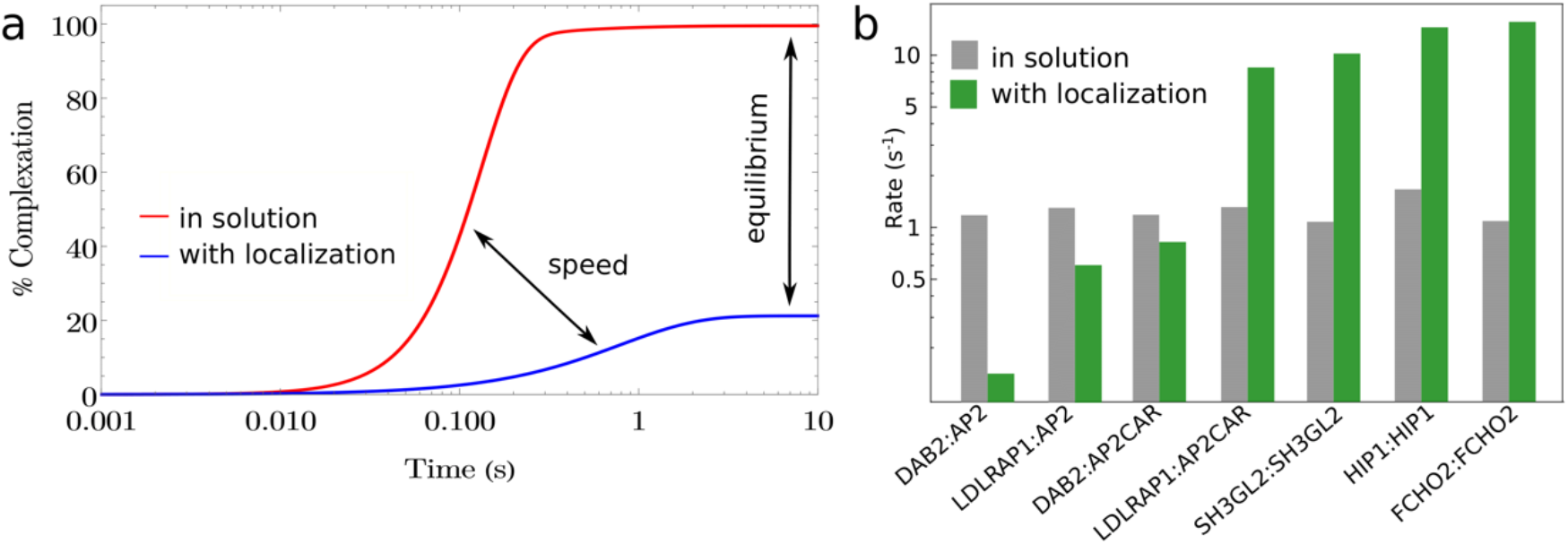
Localization to the membrane can accelerate association times for physiologic binding partners. a) For binding between the partners LDLRAP1 and AP2, both of which bind specific plasma membrane lipids, localization to the membrane accelerates the relaxation to equilibrium, in addition to dramatically shifting the equilibrium towards the complexed state. b) For some partners such as DAB2 and AP2, they have slow lipid binding and localization reduces equilibration time, but for others with moderate lipid binding rates, relaxation times are faster to the more stable equilibrium. For the AP2CAR protein, the lipid on rate is increased relative to AP2, due to binding to transmembrane cargo.

We first provide relevant background on bimolecular kinetics for two-component systems in a single volume. We derive a new localization rate to capture the influence of geometry, diffusion, and affinity on binding of a molecule from solution to a membrane. We define our model, and our target MFPT, which describes the nonequilibrium relaxation from a well-mixed unbound ensemble to the final equilibrium ensemble. Our model is sufficiently complex to study how time-scales are controlled by (i) dimensional reduction (ii) membrane ‘sticking rate’ (iii) protein-protein association rates (iv) protein concentrations, and (v) diffusion in 2D and 3D. Our derivation of an approximate MFPT is based on constructing linear systems that quantify kinetics of simplified sub-networks, with diffusion and dimension accounted for in the reaction rate constants. The theory is remarkably accurate, and produces the correct behavior in the variables (i)-(v). For equivalent concentrations of proteins, we show that our theory more accurately describes the half-time rather than the MFPT. Lastly, we use numerical simulations of both non-spatial and (stochastic) spatial reaction-diffusion implementations of the model to validate our results and our diffusion-influenced corrections to reaction rates. We discuss limitations and future extensions of this approach to variations of this model, including multi-valent self-assembly and liquid-liquid phase separations.

## II. THEORY

### II.A Background on bimolecular association in well-mixed systems

#### Relaxation kinetics

We first provide relevant background on the kinetics of association for reversible bimolecular systems, as the functional forms inform the kinetics in the coupled system. We consider two components *P*_1_ and *P*_2_ that are homogeneously distributed in the system volume and reversibly bind,

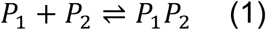

The time dependence is determined from the rate equation

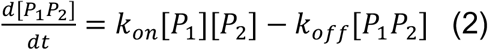

where, *k*_on_ and *k*_off_ are the macroscopic on and off rates, respectively. With the initial condition that all proteins were unbound in the solution ([*P*_1_*P*_2_]_0_ = 0), the solution to this ordinary differential equation (ODE) is well-known:

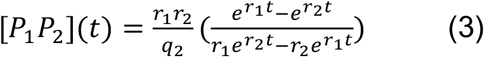

where 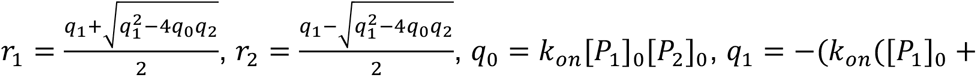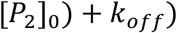, and *q*_2_ = *k*_on_. This solution is independent of the dimension of the system.

#### Characteristic timescales

A representative timescale for this relaxation process can be determined by solving for the half time *τ*_1/2_, or a mean-first passage time (MFPT). These timescales characterize the non-equilibrium relaxation from a well-defined initial state to the final equilibrium state, where the half time *τ*_1/2_ is simply when the concentration of complex *P*_1_*P*_2_ is half of the equilibrium concentration: 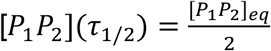. The MFPT is defined by^13^:

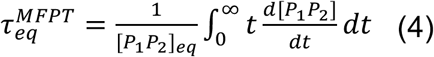

where, [*P*_1_*P*_2_]_*eq*_ is the equilibrium concentration. The analytical solution for the MFPT for Eq. (3) is quite complicated. Both exponentials in Eq. 3 are decaying with time, *r*_2_, *r*_1_ < 0 with *r*_2_ < *r*_1_ (when *r*_2_ = *r*_1_, an additional time-dependent solution is required). When initial concentrations of *P*_1_ and *P*_2_ are unequal, we find a good approximation to the MFPT is given by:

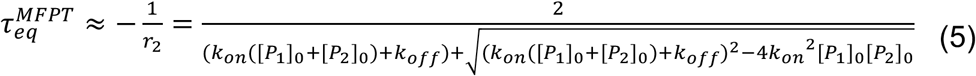

showing minor deviations with large values of 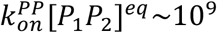. When [*P*_1_]_0_ = [*P*_2_]_0_, Eq 5 predicts the MFPT for weak to moderate binding, but for strong binding, the prediction is too fast and converges to the half-time, as we will see explicitly below.

For pseudo-unimolecular binding, [*P*_1_]_0_ ≪ [*P*_2_]_0_, the reversible MFPT simplifies to

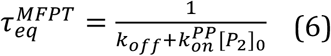

Similar to Eq 5, when [*P*_1_]_0_ = [*P*_2_]_0_, this timescale again predicts the MFPT for weak to moderate binding, but converges to the half-time for strong binding. This convergence is complete for irreversible binding, where when *k*_*off*_ = 0 and [*P*_1_]_0_ = [*P*_2_]_0_, Eq 6 is instead *equivalent* to *τ*_1/2_, not the slower MFPT. Thus, we summarize that Eq. 6 is quite accurate in describing the MFPT of reversible association for unequal concentrations of partners, and for equal concentrations, the accuracy depends on the binding strength. We find the same trend below for our coupled system MFPT.

### II.B Background on model of protein association with membrane localization

#### Model description

Now, in addition to reversible binding between our two proteins, we give each protein an additional binding site that allows it to reversibly localize to the membrane via interaction with a specific type of lipid, *M*. Thus, it is a 1:1 Langmuir binding to a surface of diffusible binding sites, not an adsorption model, allowing each protein-protein complex to potentially bind two lipids (Fig 2). We previously characterized the equilibrium behavior of this model^36^, so we briefly summarize the features here. Proteins diffuse and reversibly bind to one another both in solution (3D) and on the membrane (2D). The full set of 10 reversible interactions are illustrated in Fig 1 (see Methods Eq 18). There are nine distinct species possible: *P*_1_, *P*_2_, *M*, *MP*_1_, *P*_2_*M*, *P*_1_*P*_2_, *MP*_1_*P*_2_, *P*_1_*P*_2_*M*, and *MP*_1_*P*_2_*M*.

**Figure 2.**
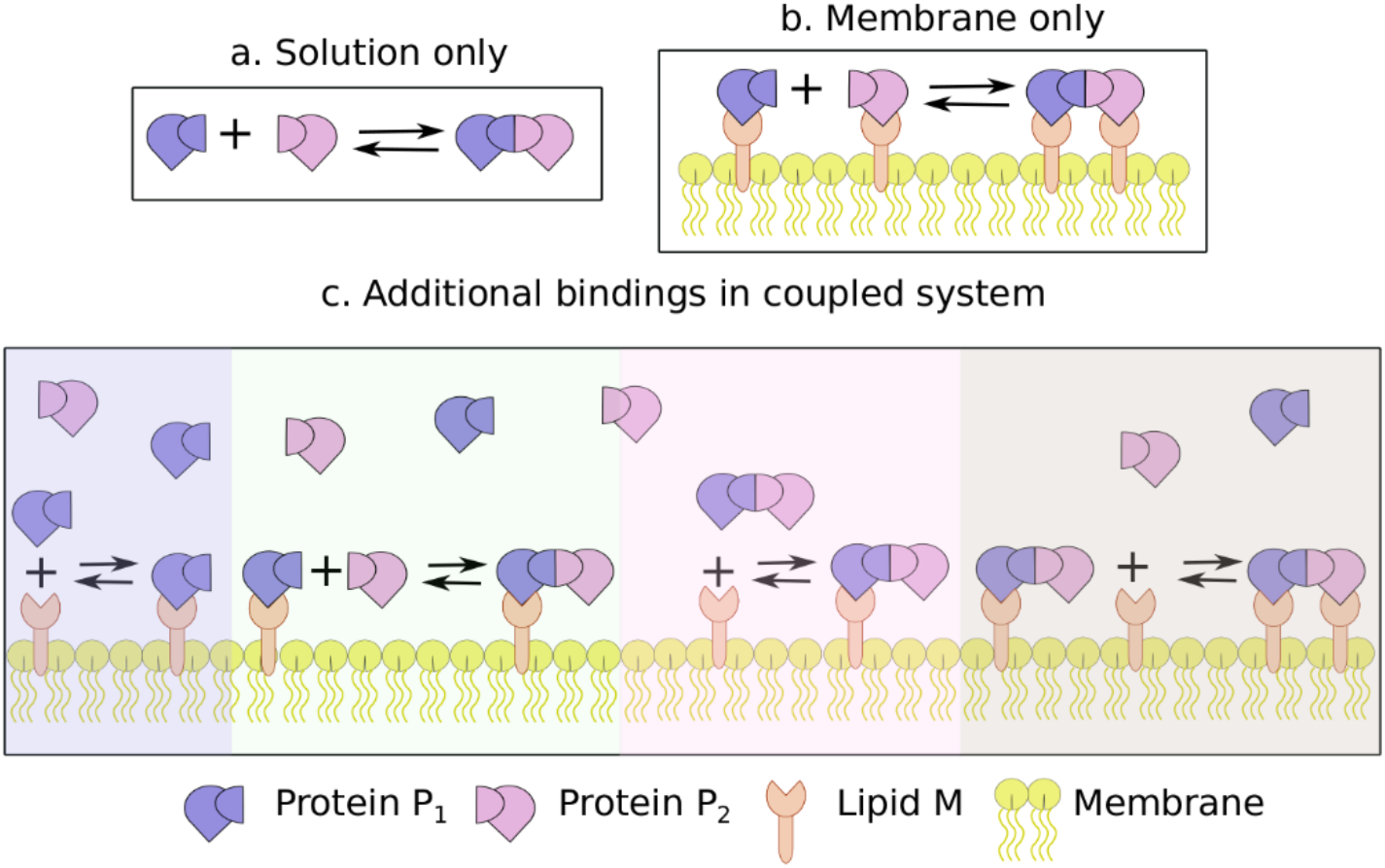
Model of bimolecular protein association with reversible membrane association: a) Two proteins *P*_1_ and *P*_2_ bind reversibly in 3*D* solution. b) Proteins are restricted to the membrane surface through binding a lipid *M* using a separate binding interface, performing a reversible 2*D* interaction. c) Coupling between the 3*D* and 2*D* domains, where the three left-hand interactions involve binding from 3*D* → 2*D* of either a single protein to a lipid, a single protein to a membrane bound protein, or a protein complex to a lipid. These are thus 3*D* searches. The final interaction on the right is purely 2*D*, where a membrane bound protein complex binds an additional lipid. For each of these 4 interactions, the pink and blue proteins can be swapped, resulting in 8 distinct reactions.

#### Macroscopic vs microscopic rates

In order to focus our theoretical analysis on the non-spatial rate equations, we incorporate the influence of diffusion and geometry on reaction rate constants. The mathematical relationships between the macroscopic rates *k*_on_ and *k*_off_ and the microscopic rates *k*_a_ and *k*_b_ are summarized in Table 1 with references. The relationships derive from the Smoluchowski model^38^ and are discussed elsewhere (e.g. see ref^27^). For numerical simulations, we primarily solve the coupled rate equations which use macroscopic rates. To accurately compare the kinetics of this non-spatial model with an explicit spatial representation using the single-particle reaction-diffusion model^39^, the macroscopic rates needed for the ODE simulations must be derived from the microscopic rates, or vice versa. The final equilibrium state obtained from both numerical approaches will be identical because of their conserved equilibrium constant, 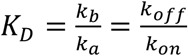, in all dimensions.

**Table 1.** Macroscopic rates with explicit diffusion dependence.

| Dimension | Rate |
| --- | --- |
| 3D<sup>40</sup> | $k_{on}^{3D} = \left( \frac{1}{k_a^{3D}} + \frac{1}{4\pi\sigma D_{tot}^{3D}} \right)^{-1}$ |
| 2D<sup>9</sup> | $k_{on}^{2D} = \left[ \frac{1}{k_a^{2D}} + \frac{1}{8\pi D_{tot}^{2D}} \left( \frac{4\log(\frac{b(\rho)}{\sigma})}{(1 - \sigma^2/b(\rho)^2)^2} - \frac{2}{1 - \sigma^2/b(\rho)^2} - 1 \right) \right]^{-1}$ |
| Localization &<br>adsorption<sup>this paper</sup> | $k_{on}^{loc} = \frac{1}{M_0 L} \left( \frac{L}{3D_{tot}} + \frac{1}{k_{on}^{PM} M_0 L} \right)^{-1}$ |

microscopic rates $k_a$ , binding radius $\sigma$ , diffusion constant $D_{tot} = D_1 + D_2$ , density $\rho = \frac{\max(N_{P1}, N_{P2})}{A}$ , surface
area $A$ , length $b(\rho) = 2\sqrt{\frac{1}{\pi\rho} + \sigma^2}$ , length $L = \frac{V}{A}$ , lipid concentration $M_0$ , 3D binding rate $k_{on}^{PM}$ .

#### 2D binding rates from 3D rates

Given a 3D association rate for a binding pair, the corresponding 2D rate will differ both in magnitude and dimensions. The macroscopic rates will explicitly account for changes in diffusivity from 3D to 2D (Table 1), hence we must specify how the microscopic association rate will transform. Localization restricts the translational and the rotational movement, limiting the orientations the two surface-bound species can sample. While the exact scale of the change thus depends on the molecular properties of the two species^41, 42^, relative scales between equilibrium constants in 3D and 2D is on the molecular (~*nm*) lengthscale^41^. Here we approximate this change as fully captured by the relative microscopic association rates, such that 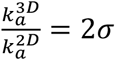, where *σ* = 1*nm*. We assume that the microscopic unbinding rate is unchanged, 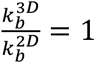. Because the relative *macroscopic* association rates may not be altered by exactly 2*σ* (see Table 1), the macroscopic off rates may not have the same value. This relation is applied to both the protein-protein association rates in 2D, and the protein-lipid association rates in 2D.

Although species on the membrane have concentration in units of particles/μm^2^ and species in solution have concentration of particles/μm^3^, for the rate-equations, all species can be solved in volume units, or equivalently, in dimensionless copy numbers, by rescaling the 2D binding constants by solution volume (V) to membrane surface area (A) ratio (V/A). Thus, we can define a (dimensionless) dimensionality factor,

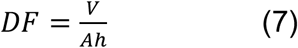

where *h* is of the order of 2*σ* and is defined by 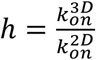. To differentiate 3D and 2D binding rates, we will now explicitly retain 2D superscripts (e.g. 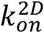) with no superscript for 3D rates.

### II.C Deriving rates of localization to a surface with geometry and diffusion correction

The rate of proteins binding to lipids is dependent on the time to diffuse to the membrane surface, and the probability of sticking to the lipid-covered surface upon arrival. In a well-mixed approximation (i.e., ODEs), the rate of lipid binding does not account for this diffusional search to the surface, as the lipids are effectively accessible throughout the solution volume. However, we can approximately correct for the impact of the diffusional search to the surface on these 3D reaction rates using the Smoluchowski model^38^ with a partially absorbing surface, or radiation boundary^18^.

We consider a density of proteins in solution *P*_0_, initially uniformly distributed in a rectangular volume of height *L*. The lipids are restricted to the bottom surface of the volume of area *A*, at a density of *ρ*_M_ (particles/*μ*m^2^). The rate of proteins binding to the lipid particles is 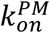. When the copy numbers of lipids on the surface is sufficiently high (*ρ*_M_*A* ≫ *P*_0_*AL*), binding of each protein to a lipid does not change *ρ*_M_ appreciably. The 1:1 binding model can thus be replaced by a surface adsorption model, with the adsorption rate 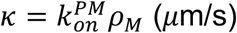. In the limit of excess lipids, the kinetics of this effectively 1D model is a very good approximation to the full 3D model with 1:1 binding 43. We are thus interested in the MFPT of a 1D diffusion model,

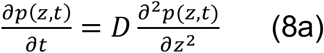

where *D* is the diffusion constant of the protein in solution. The density *p*(*z*, *t*) is subject to a radiation boundary at *z*=0 and a reflective boundary at *z*=*L* (particles do not exit the volume),

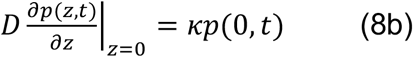

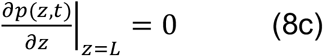

with an initially uniform distribution of particles,

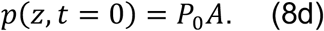

The solution to this diffusion problem and the resulting MFPT is not trivial due to the imposed boundary conditions^23^. However, previous work derived relatively simple results for this same model by solving a differential equation for the MFPT itself, giving^14^:

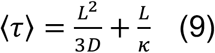

From this average timescale, we then define a new, diffusion-corrected 3D localization rate (units M^−1^s^−1^) as:

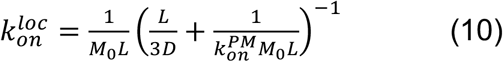

where we use the 3D concentration units for *M*_0_ = *ρ*_M_/*L*. This is one important result in our paper, because it will allow us to account for spatial contributions to relaxation times without having to solve systems of PDEs as in Eq 8. This result is also copied into Table 1 for convenience. For other non-rectangular geometries, we specify 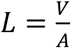. We describe the rate 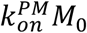 as the membrane ‘sticking rate’, as it controls the speed of protein adhesion to the lipid membrane. Our new macroscopic rate in Eq 10 thus quantifies how system length-scales, membrane sticking rate, and diffusion will impact localization events, in a similar way to the purely 3D and 2D results (Table 1). We can see that in the limit of fast diffusion and smaller length-scales *L*, 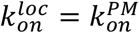, as expected. In other words, assuming lipids well-mixed in 3D is reasonable. However, this equation also shows that localization rates depend on the sticking rate, with higher values of either 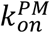 or *M*_0_ both contributing to increased sensitivity to diffusion. Overall, 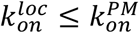.

### II.D Deriving characteristic timescales of protein association with membrane localization included

This is the primary focus of our paper, to derive an approximation to the MFPT when reversible membrane localization is included in the bimolecular association model. Now we are tracking the formation of the fully bound protein complex on the membrane surface,

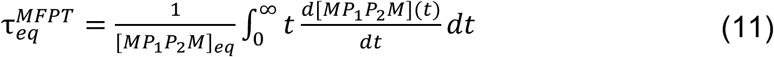

and we again assume initial conditions where both proteins and lipids are fully unbound. [*MP*_1_*P*_2_*M*]_*eq*_ is the equilibrium concentration of species *MP*_1_*P*_2_*M*, and a solution for this equilibrium value was derived in earlier work^36^, although we see below it is not needed.

#### Division of flux in two major reaction pathways

There is no closed form solution for *MP*_1_*P*_2_*M*(*t*), unlike simple bimolecular association above, due to the nonlinear couplings of now 9 species (Methods Eq 18). Even in the pseudo-first order limit where lipids and one of the proteins are in excess, the coupled linear equations are too numerous to solve for symbolic timescales. In our model, there are multiple pathways that connect our initial set of species (*P*_1_, *P*_2_, *M*) to the membrane bound complex *MP*_1_*P*_2_*M*, which we treat as the final state. We thus identify two major reaction pathways that carry flux from the initially unbound states to this final equilibrium state (Fig 3), with mathematical details provided in the Methods section. In the solution pathway, proteins first associate with one another in solution, and then localize to the membrane. We account for the 2D recruitment of the second lipid using a steady-state approximation (Eq. 22). In the membrane pathway, proteins first localize to the membrane, and then form a protein complex in 2D. In both pathways, protein-protein association is always reversible, but in the membrane pathway, the lipid binding is treated as irreversible due to subsequent stabilizing binding in 2D. We exclude from any reaction pathway the recruitment of a protein to a membrane bound protein, as it contributes negligibly to the overall reaction progression.

**Figure 3.**
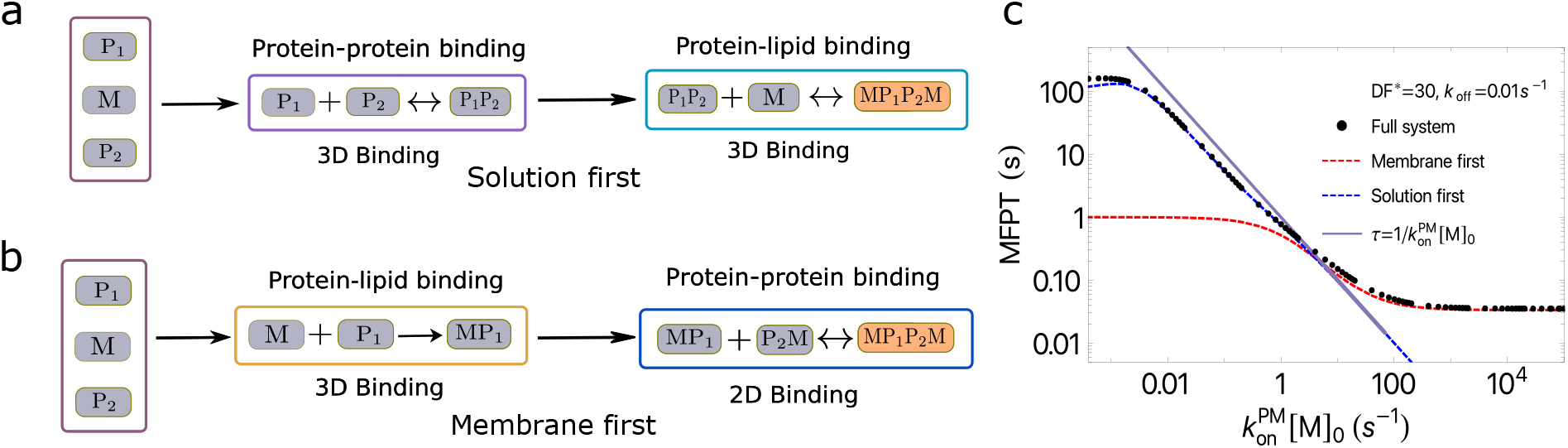
Major pathways carrying flux from initial to final equilibrium state. a) In one major pathway, reactants initially form a protein-protein complex in solution, that then localizes to the membrane. b) In the other major pathway, proteins localize to the membrane first, and then assemble a protein-protein complex in 2D. c) The solution pathway is slower, and rate-determining, when the membrane has a low sticking rate, 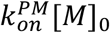, or typically when 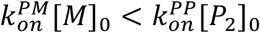. Here 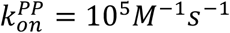, *DF*=30, [*M*]_0_ = 100*μM*, [*P*_1_]_0_ = 1*μM*, and [*P*_2_]_0_ = 10*μM*. The membrane pathway is rate-determining for all increasing membrane sticking rates. The sticking rate alone has a strong influence on the MFPT, as the gray line shows 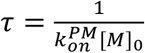, describing to a large extent the acceleration of the MFPT in the intermediate sticking rate regime.

#### Approximate expression for the MFPT

Based on our division of the relaxation kinetics into two independent pathways, we define the overall MFPT to be given by:

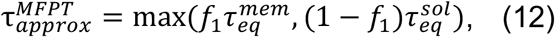

where, 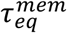 and 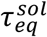are the MFPTs we will define for the membrane and solution reaction pathways, respectively. The two pathways are treated independently, but in reality they are in competition, so we must quantify the flux of proteins passing through each reaction pathway; thus we define the ratio:

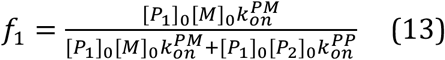

*f*_1_ quantifies the initial driving force of a protein to bind to a lipid, rather than another protein. The factor *f*_1_ is important, as it effectively defines which pathway will be taken as shown in Fig 3c. When 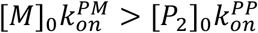, the membrane pathway is dominant, and for high membrane sticking rate, *f*_1_ → 1. Conversely, when 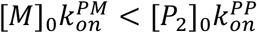, *f*_1_ drops approximately linearly with decreasing sticking rate. With the two simultaneous reaction pathways, Eq 12 chooses the *slower* reaction pathway as governing the system MFPT.

#### Assumptions for explicit MFPT results

For all the explicit MFPT expressions we make the following assumption 1) Lipids are in excess, or [*M*](*t*) ≈ [*M*]_0_, where [*M*]_0_ is the initial concentration of lipid *M*. 2) Both proteins bind to the lipids with a similar association rate, 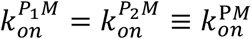. For a wide variety of physiological or *in vitro* systems, an excess of lipids relative to proteins is a good approximation, where even when only 1% of lipids act as binding partners, they still have a significantly higher copy number than most cellular proteins^36^.

#### MFPT of each pathway in the pseudo-first order limit

We will always assume that the lipid population is constant, and we further assume here one protein is in excess, [*P*_1_]_0_ ≪ [*P*_2_]_0_. Since now our binding interactions are pseudo-first order, the coupled rate equations become linear in both the solution and membrane pathways, and both the time-evolution of [*MP*_1_*P*_2_*M*](*t*) and the MFPT can be derived analytically (see Methods).

For the membrane pathway, we derive:

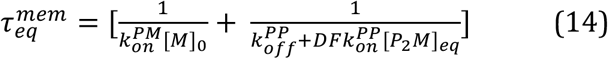

where [*P*_1_]_0_, [*P*_2_]_0_, and [*M*]_0_ are the initial concentration of proteins and lipids, 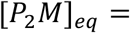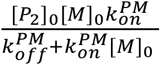 is the equilibrium concentration of species *P*_2_*M* in isolation, and the dimensionality factor *DF* (Eq 7), is defined with the ratio of protein-protein binding rates, 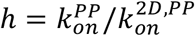. The *DF* controls how much the search space contracts on the surface, effectively concentrating the reactants, which is relative to the change in the rate from 3D to 2D, encoded in the length scale *h*. This quite simple mathematical expression is the summation of two timescales in the Eq 6 form (1) time for a free protein to irreversibly bind a lipid, and (2) time for two proteins to bind reversibly in 2D. The membrane pathway MFPT is thus dominated by the slower of the two timescales, which for lower sticking rate is 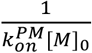, and for higher sticking rates, higher *DF*, and faster protein-protein association, transitions to the second term, 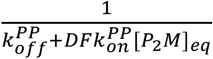.

In a similar manner, we derive the MFPT for the solution pathway to define:

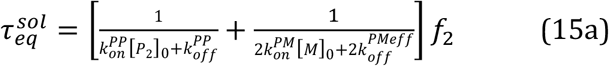

where

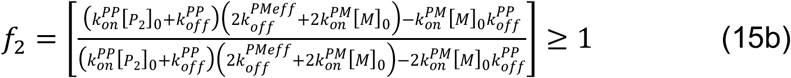

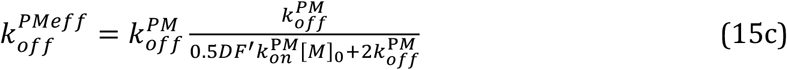

where *DF*′ is defined using the lengthscale between protein-lipid rates, 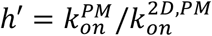. This solution pathway MFPT is essentially the summation of two times, (1) time for two free proteins to bind reversibly in solution, (2) time for a protein-protein complex to reversibly bind the membrane, via one or two lipids. The sum is scaled by a factor *f*_2_ which is 1 in the limit of low membrane sticking rate, and ≥ 1 for larger sticking rates. This scalar is derived in the Methods and results from the coupling between the protein association in solution and binding of the complex to the membrane. The dissociation of the protein-protein complex from the membrane has an effective off rate, 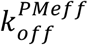, which is the only place where a dependence on the *DF* occurs. When a protein-protein complex binds to the membrane via a single lipid, it can rapidly bind a second lipid in 2D, reducing the dissociation of the complex back into solution. We made an empirical correction to this rate based on our derivation (Eq 24), so that we now see in Eq 15c that 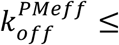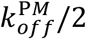. That is, the ability to bind the second lipid can only make dissociation from the membrane slower (modulo 2).

Critically, our timescales are dependent only on the initial concentrations, rate constants, and dimensionality factors of the model, with no knowledge of the equilibrium state necessary. Despite the assumptions we have made to derive these timescales (see Methods), our single expression for the MFPT:

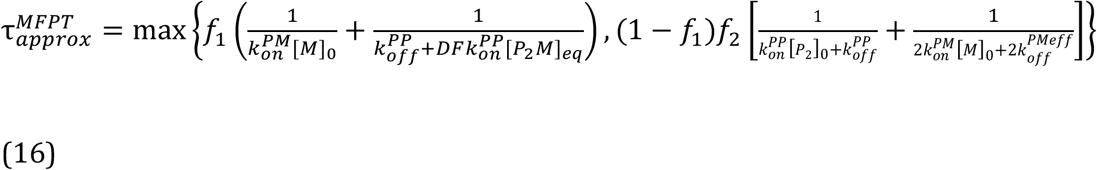

provides an excellent description of the numerical result, as shown in Fig 4. This is the primary result of our paper. As we explicitly test further below, the impact of diffusion is accounted for in the protein-protein rates (Table 1), and the impact of geometry and diffusion can be directly incorporated into the model by replacing the protein-lipid binding rate, 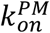, with the localization-corrected rate 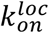 in Table 1.

**Figure 4.**
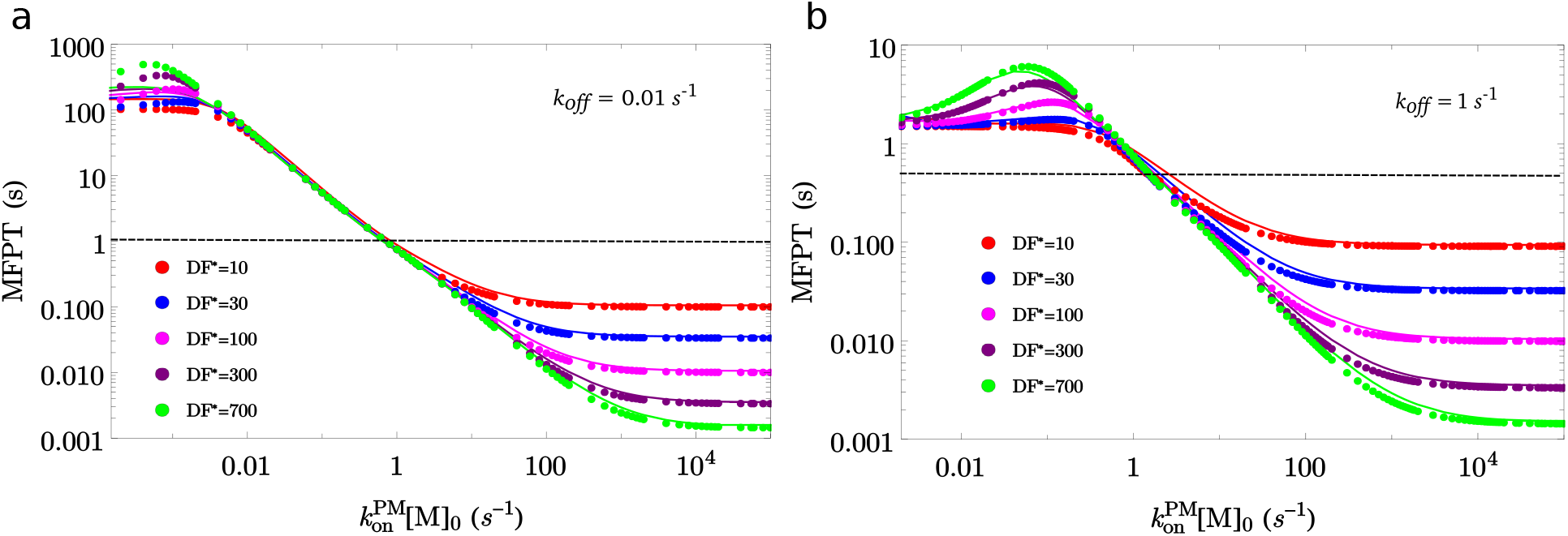
Relaxation times accelerate with increasing sticking rate to the membrane, and with higher dimensionality factor (*DF*). In the pseudo first-order limit ([*P*_2_]_0_ = 10[*P*_1_]_0_), the approximate theoretical MFPT (circles) shows excellent agreement with the numerical results from solving the full system of ODEs (continuous curves). The MFPT of the fully coupled system is plotted against the membrane sticking rate for five different values of dimensionality factor (*DF*), as indicated in the legend. We keep the copy numbers of lipids constant at [*M*]_0_ = 100*μM*, and vary 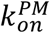 on the x-axis. The black dashed line is the MFPT for a purely solution reaction, showing that for low sticking rates, membrane localization does slow the equilibration. a) *k*_off_ = 0.01s^−1^ in all reactions. b) *k*_off_ = 1*s*^−1^ in all reactions. For both panels, 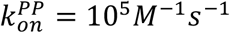, [*P*_1_]_0_ = 1*μM*, [*P*_2_]_0_ = 10*μM*.

## III: RESULTS

### III.A MFPT in pseudo first-order limit

In Fig 4 we observe excellent agreement between our theoretical predictions and the numerically calculated MFPT. Our results show how either by varying the binding rate between proteins to lipids (here from 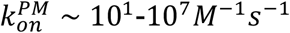), or by increasing the *DF*, one can accelerate the association times by orders-of-magnitude, from seconds to milliseconds. The MFPT is broadly divided into two regimes relative to the solution binding time-scale, with the boundaries of these two regimes essentially determined by the criteria that *f*_1_ ≈ 0.5, or 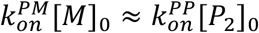.

In the slow membrane binding regime, where 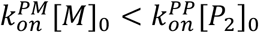, a larger fraction of proteins binds first in solution and then localize to the membrane surface, making the solution binding pathway (usually) the dominant reaction pathway (Eq. 15). Simplifying the result in Eq 15 multiplied by the factor 1 − *f*_1_, we see that for very slow binding 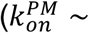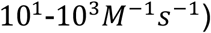 we get,

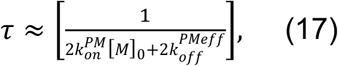

where the timescales are dominated by the time for a protein-protein complex to reversibly bind the membrane, via one or two lipids.

In the fast lipid binding regime, where 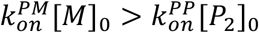, a larger fraction of proteins first localizes to the membrane and then bind one another in 2D, following the membrane pathway. In this regime, the timescales continue to accelerate, but at an increasing slower rate, until they reach a plateau. One can see this directly from Eq 14, where the prefactor *f*_1_ approaches 1, and thus the MFPT is first dominated by 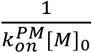. With increasing membrane stickiness, however, we switch to fully 2D binding, which is controlled by the 2D binding rate of 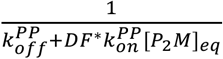. Thus the maximum time-scale is the purely 2D system relaxation time.

The membrane sticking rate has a dominant control over the MFPT in the transition regions, where *f*_1_~0.5 and both pathways will carry flux to the equilibrium state. While this is more obvious for the membrane pathway, for the solution pathway as well, this membrane sticking rate is a dominant contribution to the MFPT via the power law 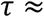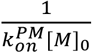. In Fig 4a for example, the slope is approximately linear on a log-log scale in the range from 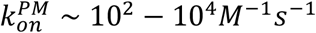. While it is not exact (Fig 3c, grey line), we nonetheless see that the MFPT in this regime is primarily determined only by the initial concentration of lipids and the protein-lipid binding rate, and not on the *DF*, the protein-protein binding rates, or the protein concentrations.

#### Role of dimensionality factor, DF

An increase in *DF* can be achieved by keeping the volume fixed and reducing the size of the membrane area. By keeping the copy numbers of proteins and lipids constant, this results in a smaller search space on the membrane, with a higher density of lipids. For moderate values of 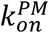, the *DF* has no impact on timescales (Fig 4). This is where localization to the membrane is rate-limiting, not the 2D binding events that can exploit reduced dimensionality. For fast sticking rates, larger *DF* drives faster timescales. This is because, as we note above, the 2D binding of proteins with one another becomes the rate-limiting step, and it is strongly influenced by the *DF*, 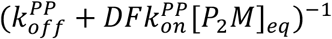, where [*P*_2_*M*]_*eq*_ depends on the membrane sticking rate directly (see Eq 14). In contrast, for slow sticking rates, we see an inverse effect, where a higher DF drives slower timescales. This is because the timescales are dominated by dissociation rates, and dissociation from the membrane is slower with higher *DF* due to the ease of binding more lipids in 2D.

#### Dependence of MFPT on the protein-protein binding rate 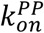

The effect of protein-protein binding strength on timescales strongly depends on whether the proteins are free in solution, or on the membrane. This is most easily seen by comparing the timescales of a purely 3D system and a purely 2D system (Fig. 5). As long as *DF* > 1, proteins in 2D exploit dimensional reduction^36^, meaning their concentration has increased relative to 3D; association kinetics in any dimension are strongly dependent on concentration. Hence, we see that despite the fact that diffusion on the membrane surface is 100 times slower than diffusion in solution, the 2D system is usually faster than the 3D system. Only when the binding rate becomes strongly diffusion-influenced (in this system at 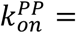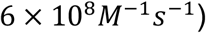 does the 2D system become slower.

**Figure 5.**
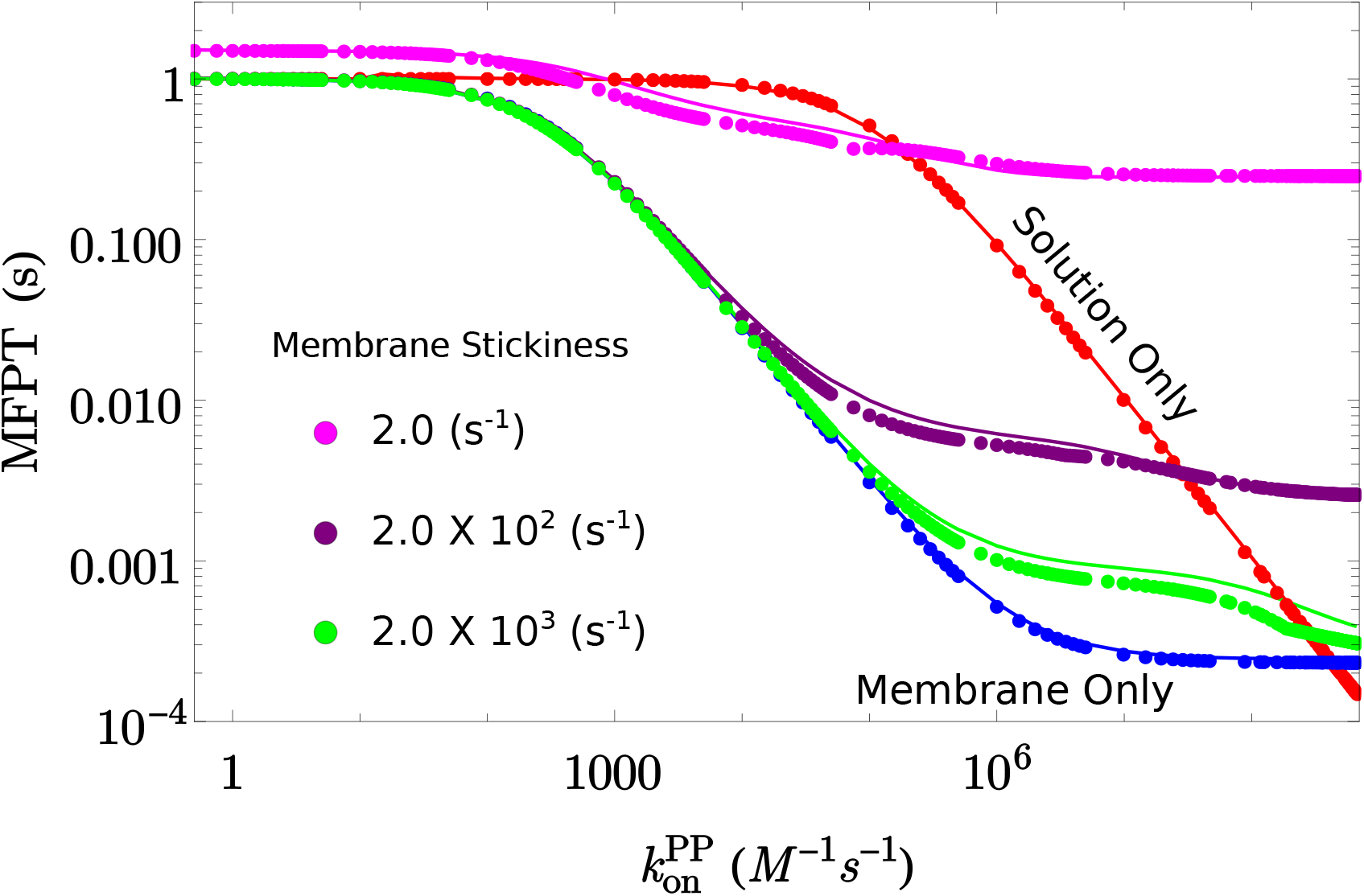
Localization to the membrane can accelerate relaxation even for moderately strong protein-protein interactions. In the pseudo-first order limit, our theoretical MFPT (circles) shows good agreement with the ODE numerical solutions (continuous curve). For the purely 3D (red) and purely 2D (blue), the simulation and well-known theory (Eq 6) are in very close agreement, as expected. For the coupled system, we show results for three values of membrane stickiness, 2, 200, and 2000s^−1^. Our theory slightly deviates from the numerical solutions in the transition regime from membrane pathway to solution pathway. The V/A ratio of this system is 0.70*μm*, and *DF* = 350, which is necessary to directly compare even a purely 3D vs 2D system. We use this value to convert a 2D rate to a 3D rate. Our macroscopic rates account for the change in diffusion from 3D to 2D, where we assume the intrinsic rates are conserved (see Table 1). *D^3D^* = 50*μm*^2^/*s*, *D*^2D^ = 0.5*μm*^2^/*s*. [*P*_1_]_0_ = 1*μM*, [*P*_2_]_0_ = 10*μM*, [*M*]_0_ = 100*μM*.

In the coupled system, we again see very good agreement (Fig 5) and observe that localization can again accelerate time-scales by up to ~1000 fold. The major difference in the coupled solution is that any speed-ups relative to the purely 3D system depend on the membrane sticking rate. For high sticking rates (200-2000 s^−1^), the coupled system is almost as fast as the purely 2D system, particularly for weaker protein association, and shows significant speed-ups relative to the 3D system, due to the rapid transition to the membrane surface. Similar to the purely 2D system, we again see a cross-over at high 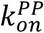 values, where it is faster to associate in purely 3D. At high values of 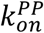, the MFPT switches to the solution pathway and essentially plateaus because the relaxation time is dominated by the time to localize to the membrane. Hence this plateau value is shifted up as the membrane sticking rate slows. In summary, for weak to moderate 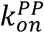, the MFPT follows the membrane pathway, and is thus rate-limited by localization and 2D protein-protein association times, whereas at higher 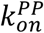, the membrane localization time becomes rate limiting.

### III.B Characteristic times for equal reactant populations [*P*_1_]_0_ ≈ [*P*_2_]_0_

Here, we will relax the assumption from section III.A where one protein had to be in significant excess of the other, such that now [*P*_1_]_0_ = [*P*_2_]_0_. The lipids are still in excess of either protein. In this case, the bimolecular association of proteins cannot be approximated as a pseudo-first order, linear set of ODEs. The rate equations are thus nonlinear in several of the unknowns. Here, we will therefore simply test our approximate MFPT derived in the pseudo-first order limit, Eq. 16. This is motivated by the relatively good agreement in the simpler two-component system (Eq. 6) between the pseudo-first order results and the equal reactant case, and the expected switch to the half-time.

In Figure 6a we compare our prediction against the numerical MFPT, finding good agreement for weak protein-protein binding, but an overly fast predicted relaxation time at stronger binding. However, this is visibly true even for the purely 3D and 2D system, where we used as the theory the approximate result in Eq 5. Thus in Fig 6b, and motivated by the behavior in the two-component system, we compare the same theoretical prediction with the numerically calculated half-time. Now we see the opposite trend, where for weak 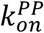, the agreement is worse, but for fast 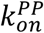, the theory does a good job of predicting the half-time. Thus the trend is the same as what happens in the two-component system, as anticipated. For strong binding, it makes sense that our prediction agrees with the half-time, because this regime is most similar to irreversible association, where the formula for the MFPT in Eq 6 becomes instead *equivalent* to the half-time for equal reactants. In summary, when reactants are in equal concentrations and protein-protein binding is moderate to strong, as we test further below, our predicted time-scale should be compared with the half-time, which is typically 1.5-2.5 times faster than the MFPT.

**Figure 6.**
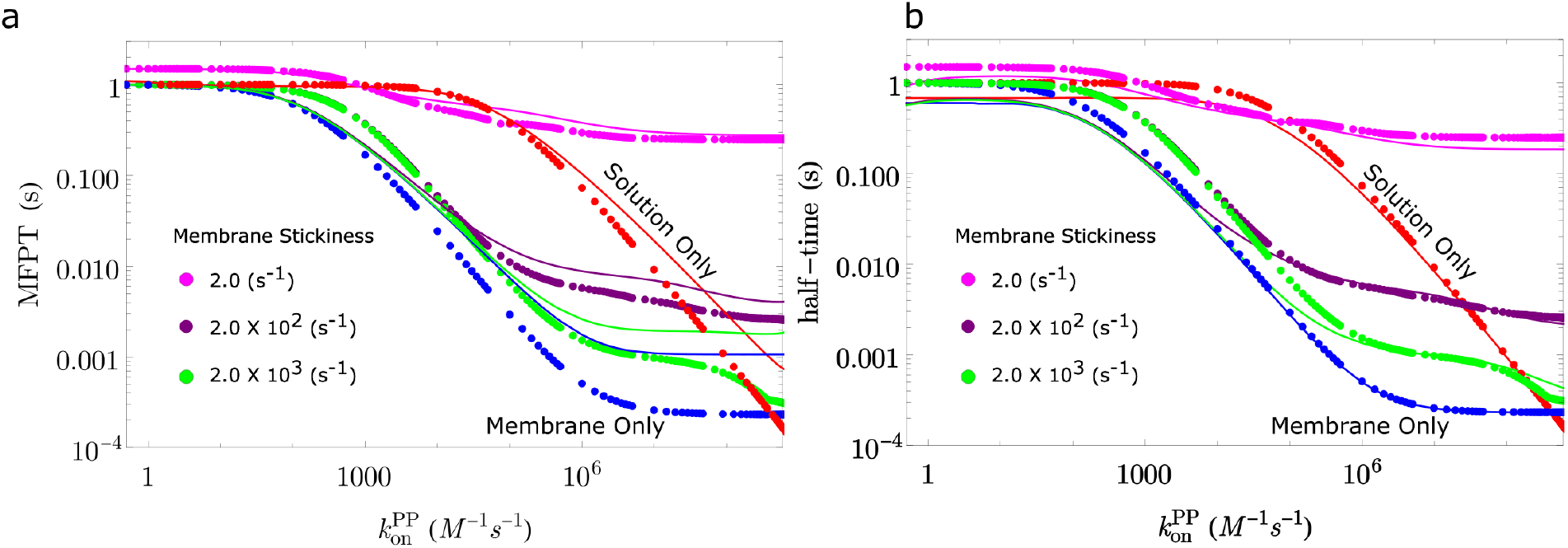
With equal protein populations, the theory is a better predictor of the half-time rather than the MFPT for moderate to strong binding. a) The MFPT of the fully coupled system is plotted against the protein-protein binding rate, for three values of membrane sticking rate. Theory in dots is in close agreement with numerical results (solid curves) for weak binding, but shows deviations at stronger binding even for pure 3D (red) or pure 2D (blue) binding. b) The theory here is the same, but the numerical results (solid curves) now report the half-time to reach equilibrium. The agreement is closer as binding strength increases. *D*^3D^ = 50*μm*^2^/*s*, *D*^2D^ = 0.5*μm*^2^/*s*. [*P*_1_]_0_ = [*P*_2_]_0_ = 10*μM*, [*M*]_0_ = 100*μM*.

### III.C Effect of diffusion and spatial geometry can be captured in rate-constants to predict the MFPT

#### Accounting for diffusion in non-spatial rate equations

Our results above are based on deriving approximate MFPTs for the system of coupled rate equations, with comparison against numerical solutions solved using non-spatial ODEs. The effect of explicit spatial representations and diffusion on the system dynamics is two-fold. First, even for a well-mixed system purely in 3D or 2D, partners must diffuse to contact with one another, which directly impacts the bulk or macroscopic rates of association, *k*on. Second, we have two domains in our system, and the membrane domain has to be arrived at before any protein-lipid interactions can occur. In other words, the lipids are not well-mixed in a real system, as is assumed in the ODEs. For the first point, we use the rates in the first two rows of Table 1, which in 3D are accurate and in 2D are very good estimators for the kinetics. For the second point, we use our localization corrected rate derived in Section II.D, 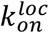 (Table 1), which adjusts the protein-lipid binding rate, 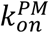 based on the height of the system, the sticking rate to the membrane, and the diffusion constant. In Fig 7, we see that this correction does a remarkably good job in improving agreement between the ODE solution and the explicit RD solution.

**Figure 7.**
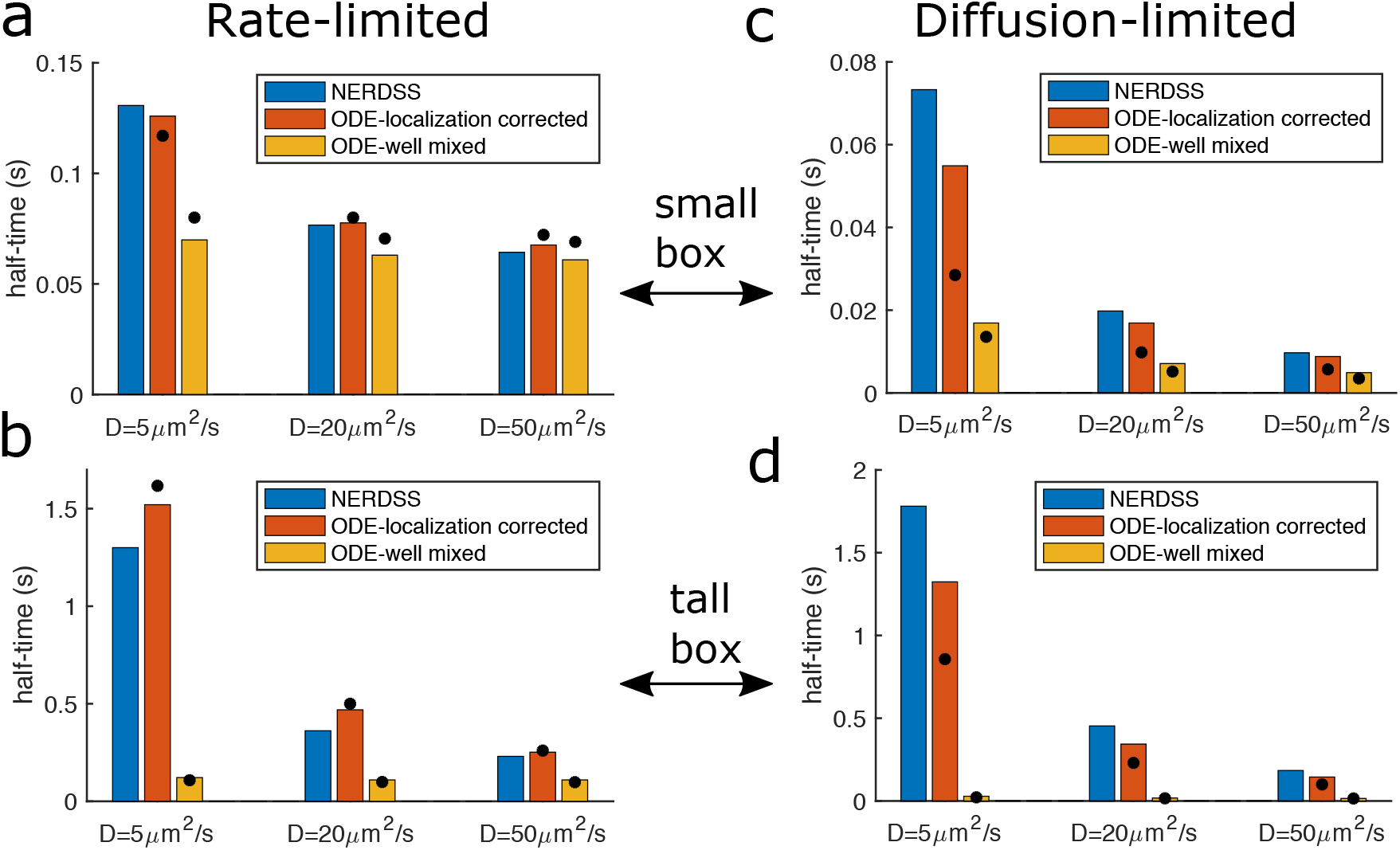
Our model predictions are in good agreement with reaction-diffusion simulations that quantify the impact of explicit spatial dynamics on time-scales. a) For a more rate-limited reaction, diffusion has quite modest impact on the half-time. In all panels, diffusion slows from [*D*^3D^, *D*^2D^] of [50,0.5] to [5,0.05] *μ*m^2^/s. The height of the small box is 0.75 *μ*m. The RD simulations (blue bars) slow the most, due to the time required to localize to the membrane, and we see the ODEs solved with localization-corrected rates, where we replace 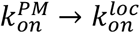 orange bars) provides very good agreement with the explicit spatial simulations. ODEs without localization correction in yellow bars. Theoretical predictions for both sets of rates in black circles. b) With the same rates but a taller box height of 5 *μ*m, simulation times are slower and the RD simulations are more sensitive to diffusion, as expected, with again very good agreement using 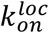. c) For a diffusion-limited reaction, even in the small box the half-time is more sensitive to diffusion, for all three models. d) Diffusion-limited reaction in the tall box is most sensitive to diffusional slow-downs. Here, changing diffusion by a factor of 10 does change the half-time by about a factor of 10 as well, which is a significant change from panel (a). In all systems, [*P*_1_]_0_ = [*P*_2_]_0_ = 1*μM*. The lipid copy numbers are fixed at 17000/*μ*m^2^, and the membrane surface area is 0.2209 *μ*m^2^. In the small box, [*M*]_0_ = 37.55*μM*, and in the tall box [*M*]_0_ = 5.64*μM*.

#### Rate-limited reaction regime

In the more rate-limited regime, where 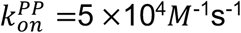 and, 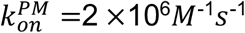, we see that the impact of diffusion on the *well-mixed* kinetics is negligible, as despite dropping from *D*^3D^/*D*^2D^= 50/0.5 *to* 5/0.05 *μm*^2^/*s*, the ODE solution shows minimal slow-downs (yellow bars in Fig 7a-b). We note that we do account for diffusion in the ODEs here—all systems have the same microscopic rates, but the macroscopic rates become slightly smaller with slower diffusion. The RD timescales slow a bit with slower diffusion. This is due to the time to reach the membrane before binding lipids, and is thus significantly more dramatic going from a system height of 0.75*μ*m (Fig 7a) to 5*μ*m (Fig 7b). By correcting for this rate of lipid binding from solution (orange bars in Fig 7), we see excellent improvement in describing the RD MFPT in both system sizes for this more rate-limited reaction. We note that with the taller box, the protein concentrations are the same, and thus the lipids are not in as significant excess of the proteins. For all systems, the theoretical time-scale is determined by following the membrane pathway.

#### Diffusion-limited reaction regime

In the more diffusion-limited regime, with microscopic rates of 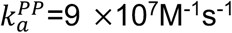 and 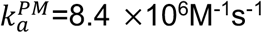, even the non-spatial ODE results show slow-downs with diffusion (Fig 7c-d). For the small box (Fig 7c) the timescales are more sensitive in the RD simulations, as expected, with a decrease of about 6x from the fast to slow diffusion regime. In this diffusion-limited system, the localization corrected ODE solution (orange bars) is not as accurate at reproducing the RD results. When comparing the time-dependent kinetics, we see that the ODE does accurately describe short time kinetics, but at longer times, the RD systems decay more slowly to equilibrium, causing a deviation in the half-times. This is perhaps not surprising, as when distinct proteins have equivalent starting concentrations, as we use here, even irreversible association is known to decay more slowly than rate equations predict^44, 45^. This is because as the reaction proceeds, regions of isolated A and B molecules develop^46^, which will cause a slow-down in the relaxation. These deviations are detectable only in the more strongly diffusion-influenced regime. For the taller box, the system is even more sensitive to diffusion, as expected, decreasing ~10 × from fast to slow diffusion, and showing the largest mismatch with the well-mixed ODEs. For this system, the predicted half-time is in the transition region between being dominated by the solution vs membrane pathway. For the localization-corrected rates, the theory always follows the solution pathway, and although the predicted timescales are a bit too fast as diffusion slows, they capture the order-of-magnitude slow down relative to the well-mixed model.

## IV. Discussion and Conclusions

We have shown here how the time-scales of bimolecular association between protein populations can be dramatically shifted by localization to the membrane surface. The degree to which this assembly speed is accelerated or slowed is strongly dependent on the relative sticking rate to the membrane. For fast sticking rate, proteins will exploit the lower dimension of the surface to assemble more quickly. For slower sticking rate, protein assembly is rate-limited by localization to the surface, which often reduces speeds relative to pure 3D assembly. Critically, we address here how the geometry of the system and diffusion will potentially slow this sticking rate. By accounting for these spatial effects directly in our rate constants, we can retain a relatively simple functional form for our predicted MFPT, as we derived from the non-spatial rate equations. One can then directly assess whether diffusion or geometric changes to the system will impact the overall MFPT, as illustrated in our Fig 7 examples. Due to the importance of reaching the membrane, we find that the height of the volume typically has a bigger impact on the MFPT than the diffusion constant does. Perhaps counterintuitively, the speed of diffusion on the membrane surface is rarely a rate-limiting factor, where as long as the *DF* is >1, the increased concentration on the surface accelerates 2D assembly.

We derive the MFPT for our multi-component model based on assuming lipids and one binding partner are in excess, and that both proteins bind the lipids with similar affinities. Thus, although we see excellent agreement for variations in all model parameters, including population, kinetic, and spatial factors, the main MFPT result (Eq 16) would have to be adapted if one protein bound lipids significantly weaker than the other, or if lipids were in short supply. In many physiologic membranes, lipid abundance will exceed most proteins^36^, but for proteins that target a small patch of membrane for assembly, for example, the lipids might be in limited supply. Further, although we are able to account for the lengthscale of a cell system via our localization rate, this assumes a simple geometry such as a box or sphere. For more complex cell geometries of either the membrane or the volume, a more explicit treatment of the diffusional search may be necessary. Last, for equivalent protein concentrations, [*P*_1_]_0_ = [*P*_2_]_0_, we find that our theory still applies quite well, but only when the time-scale is interpreted as the half-time, rather than the slower MFPT. This is not particularly surprising given this same behavior for the two-component system in a single volume domain. Perhaps the biggest room for improvement is more precisely capturing the behavior of the explicit reaction-diffusion simulations^24^, which tend to be slower for the strongly diffusion limited regime than our prediction, although in good agreement for more rate-limited reactions (Fig 7). This would likely require adding the spatial components more explicitly into the model.

By allowing us to predict how protein association times are controlled by both solution and membrane properties of a system, our model provides a useful foundation to anticipate assembly speeds in more complex systems. Our model lacks the multi-valent interactions that stabilize higher-order cluster formation and self-assembly, but it has key parallels with phase separated systems which are prevalent in cell biology^47, 48^. Our model contains two domains which create a dilute (solution) and a dense (membrane-bound) phase of identical components, and there is no barrier to transitioning between these phases. Within each phase, both the stability and the kinetics of the protein-protein interactions are altered. Hence our model creates a steady-state solution where the densities have changed due to the capacity of all components to stick to the surface. The relative sticking rate to the surface and the *DF* are thus key in controlling both the speed of formation and ultimately the steady state populations of both phases. In biology, some liquid-like droplets do seem to exploit membrane localization to drive dense phase assembly49, 50 and the theory developed here, while simpler, provides insight into the kinetics of these processes.

The kinetics of bimolecular association is also a key step in structured self-assembly^51^, where nucleation times must generally be slower than growth speeds to ensure productive assemblies^52^. Theoretical models have predicted how binding rates and concentration can control self-assembly in solution^53, 54^, as well as the origins of the lag-time^55^. Our model provides independent variables—including the membrane sticking rate and *DF*—that can help tune the kinetics of the protein association by creating a dense and a dilute phase at a particular spatial location. While modeling self-assembly in and out of equilibrium ultimately depends on inherently spatial and structural details^24, 56^, kinetic models can be effective tools for studying assembly kinetics^57^ even without spatial resolution^58^. Exploiting the membrane surface as we show here represents a powerful dimension for improving and understanding control of association and higher-order assembly processes.

## V. Methods

### V.A Coupled rate equations for the full system

The time-dependence of bimolecular association between protein partners P1 and P2, when they can additionally localize to membrane lipids *M* is given by the following coupled differential equations in the well-mixed limit:

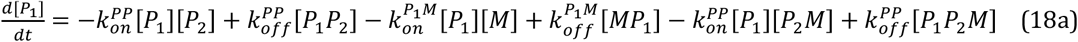

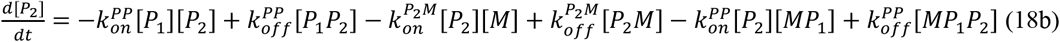

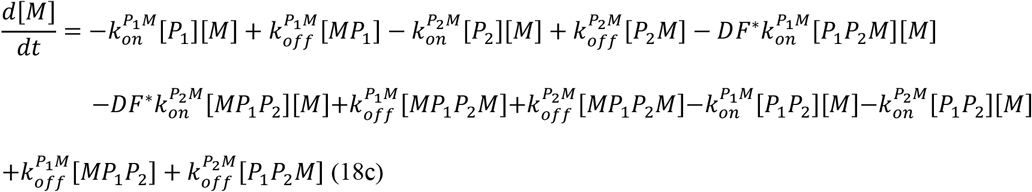

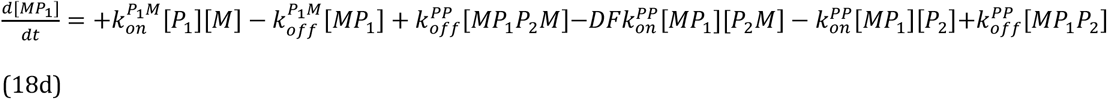

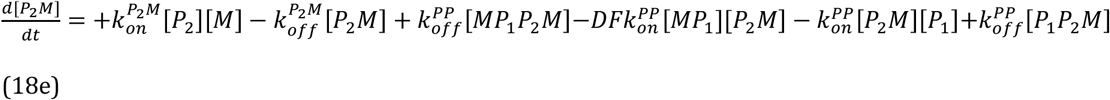

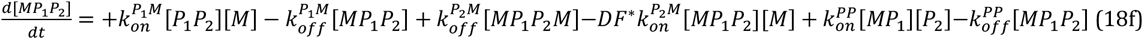

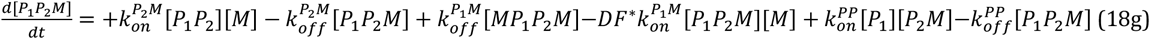

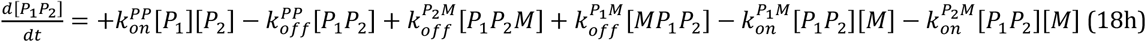

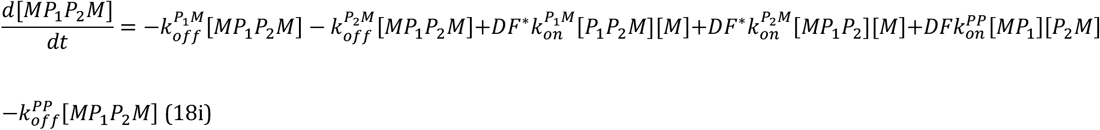

In general, the protein-lipid binding rate depends on individual binding partners and changes from one binding pair to other. However, for simplicity, we assume a similar association rate for both *P*_1_ and *P*_2_, 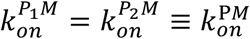.

### V.B Division of flux into dominant reaction pathways

Instead of solving the rate equations for the full system above, we simplify by identifying two major reaction pathways. The MFPT are nonetheless mechanistic, although we make minor empirical corrections to the solution pathway to improve agreement.

#### Membrane first reaction pathway

In the membrane first reaction pathway, we solve the following set of differential equations,

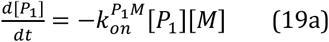

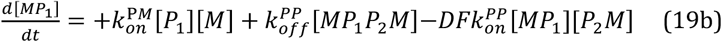

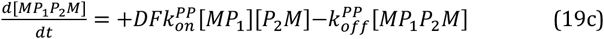

using pseudo first-order approximations, such that [*M*](*t*) = [*M*]_0_ and from the original model, we assumed [*P*_1_*P*_2_*M*] = [*MP*_1_*P*_2_] = [*P*_1_*P*_2_] = 0. Also, 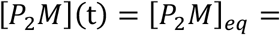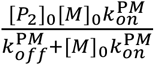, where this definition of proteins *P*_2_ on the membrane is derived assuming an isolated pairwise equilibrium between protein and lipids. We further assume irreversible lipid binding, 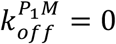, because the formation of *P*_1_*M* results in its participation in additional binding reactions on the surface, which largely prevents dissociation back to solution. We thus have a linear system of our unknowns, which can be solved exactly via the eigenvalues and eigenvectors. The system has two non-zero eigenvalues 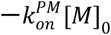 and 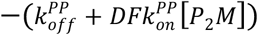, resulting in the time-dependent solution:

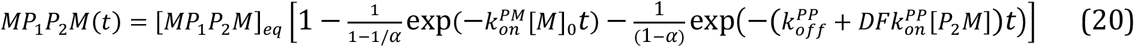

where 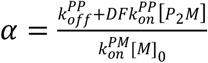. The MFPT of this membrane pathway is the negative sum of the inverse eigenvalues, or Eq 14 of the main text.

#### Solution first reaction pathway

In the solution first reaction pathway, we will solve the following set of differential equations:

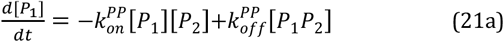

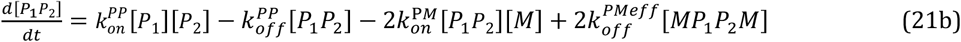

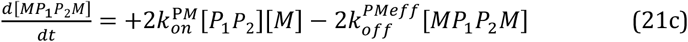

using similar assumptions as before, where [*M*](*t*) = [*M*]_0_, and from the original model, we assumed [*P*_2_*M*] = [*MP*_1_] = 0. We further assume a rapid equilibrium between a protein complex on the membrane with one (*P*_1_*P*_2_*M*) versus two lipids (*MP*_1_*P*_2_*M*) bound, because the second lipid binding reaction takes place in 2D with an abundant lipid population. We use this assumption to derive an expression for the effective off rate of the protein-protein complex to dissociate from the membrane. In particular, at equilibrium we have that,

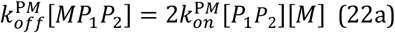

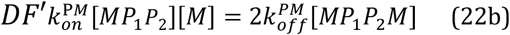

where factors of 2 emerge due to having two sites on a protein-protein complex to bind a lipid (*P*_1_ and *P*_2_). *DF* is the dimensionless dimensionality factor (*DF*′ = *V*/*Ah*′). We are looking for the effective rate that will give us:

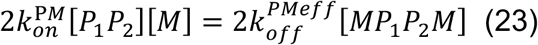

which is 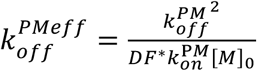. We now make an empirical correction, and will instead use:

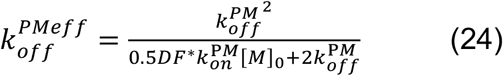

This correction is used because for low sticking rate, the dissociation rate should not exceed the rate for unbinding a single lipid. Overall, this expression states that the dissociation rate for a protein-protein complex from the membrane is typically much slower than for unbinding a single lipid, because we are effectively accounting for the dissociation of up to two lipids.

With this linear system, we again have two non-zero eigenvalues, 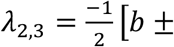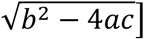, where *a* = 1, 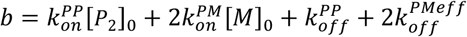 and 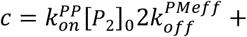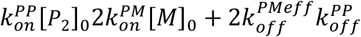. The MFPT is the sum of their inverses, which simplifies to:

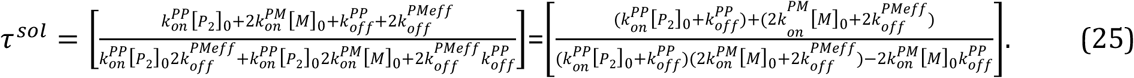

This expression is dominated by two timescales 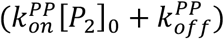 for protein-protein binding in solution, and 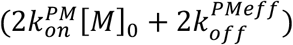 for binding and unbinding of the protein-protein complex from the membrane. In the limit of low membrane sticking rate (small 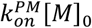), this expression simplifies to

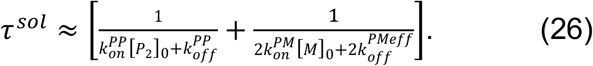

Empirically, this timescale is a bit too fast for larger sticking rates, whereas the previous prediction is a bit too slow. If we take the average, we finally arrive at Eq. 15 of the main text. The Eq. 15 result overall has the same limits as Eq 25 (*f*_2_ → 1 with 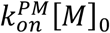 small) but with better agreement in the transition region where the solution and membrane pathways have comparable time-scales.

### V.C Simulation details

We solve the non-spatial coupled ODEs numerically in Fortran by the standard Runge-Kutta integration scheme until the steady state was attained with an initial condition that all the proteins and lipids were unbound, and all proteins were in solution. First, we derive the time dependent concentration of the fully bounded complex *MP*_1_*P*_2_*M*(*t*). Next, we derive the derivative of *MP*_1_*P*_2_*M*(*t*) using the finite difference method and multiply the derivative with time. Then for the MFPT, we integrate the product using numerical integration with the midpoint rule. For ODE simulations the default value of 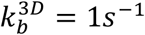, unless it is specifically stated. For numerical derivations, we first derive 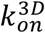 from the first equation of Table 1, then 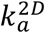 from 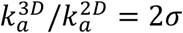, and then we derive 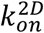 from the second equation of Table 1. For all models, we used *σ* = 1 nm.

For the single particle reaction-diffusion (RD) spatial simulations, all parameters are provided in Supplemental Table 1. To summarize, simulations were performed using the NERDSS software^24^, in a rectangular volume with the membrane defined as the bottom surface, and reflecting boundaries on the other ‘walls’. Simulations were performed with an implicit lipid model^43^, after verifying the kinetics were identical to the explicit lipid model. Kinetics were collected from 30-48 trajectories. The MFPT was calculated in MATLAB using numerical integration with the midpoint rule. Diffusion coefficients in 2D were 100 times slower than the specified 3D value.

## Acknowledgments

This work was supported by NIH MIRA grant R35GM133644 to MEJ, and used resources from the NSF XSEDE Stampede2 supercomputer through XRAC grant MCB150059.

